# Validity and reliability of extrastriatal [^11^C]raclopride binding quantification in the living human brain

**DOI:** 10.1101/600080

**Authors:** Jonas E Svensson, Martin Schain, Pontus Plavén-Sigray, Simon Cervenka, Mikael Tiger, Magdalena Nord, Christer Halldin, Lars Farde, Johan Lundberg

## Abstract

[^11^C]raclopride is a well established PET tracer for the quantification of dopamine 2/3 receptors (D_2/3_R) in the striatum. Outside of the striatum the receptor density is up to two orders of magnitude lower. In contrast to striatal binding, the characteristics of extrastriatal [^11^C]raclopride binding quantification has not been thoroughly described. Still, binding data for e.g., neocortex is frequently reported in the scientific literature. Here we evaluate the validity and reliability of extrastriatal [^11^C]raclopride binding quantification. Two sets of healthy control subjects were examined with HRRT and [^11^C]raclopride: i) To assess the validity of extrastriatal [^11^C]raclopride binding estimates, eleven subjects were examined at baseline and after dosing with quetiapine, a D_2/3_R antagonist. ii) To assess test-retest repeatability, nine subjects were examined twice. Non displaceable binding potential (*BP_ND_*) was quantified using the simplified reference tissue model. Quetiapine dosing was associated with decrease in [^11^C]raclopride *BP_ND_* in temporal cortex (18±17% occupancy) and thalamus (20±17%), but not in frontal cortex. Extrastriatal occupancy was lower than in putamen (51±4%). The mean absolute variation was 4-7% in the striatal regions, 17% in thalamus, and 13-59% in cortical regions. Our data indicate that [^11^C]raclopride PET is not a suitable tool for D_2/3_R binding quantification in extrastriatal regions.

## Introduction

The dopamine (DA) system is of key interest both in normal brain function and in the pathophysiology of neurological^1^, and psychiatric^2,3^ disorders. Striatum is the brain region with the highest concentration of dopamine receptors^4^ and also the most studied using positron emission tomography (PET). In recent years, quantification of dopamine receptors in extrastriatal regions has received more interest^2,5^. Specifically, striatal and extrastriatal availability of the dopamine D2 receptor family has been of particular interest in psychiatry research as drugs targeting D_2/3_ receptors (D_2/3_R) is an established treatment of psychosis and mood disorders^6^.

The dopamine D2/3R radioligand [^11^C]raclopride was developed in the 80’s^7^ and is one of the most frequently used PET radioligands to date. Due to its relatively low affinity to D_2/3_R (Kd = 1.3 nM) [^11^C]raclopride has primarily been used to study receptor availability in striatal regions. Extrastriatally, the concentration of D_2/3_R is up to two orders of magnitude lower than in striatum ^8^. To study regions with low levels of D_2/3R_, high affinity radioligands have been developed, e.g. [^11^C]FLB-457 (Kd = 0.02 nM) and [^18^F]fallypride (Kd=0.03 nM)^9,10^. These tracers are, however, not ideally suited to quantify D_2/3_R in striatum. If [^11^C]raclopride binding to extrastriatal D_2/3_R could be shown to be validly and reliably quantifiable, fewer PET-examinations would be required for studies where D_2/3_R in the whole brain is of interest. Although there is some indication of reliable quantification of the extrastriatal [^11^C]raclopride signal^11,12^ (i.e., adequate test-retest properties), there is a lack of data supporting quantifiable specific binding in these regions. In spite of this, several PET-laboratories, including our own, have applied [^11^C]raclopride to measure extrastriatal D_2/3_R availability in thalamus^13–15^, and in cortical regions^16–18^.

In a statistical context, reliability is the repeatability or consistency of a measurement. In PET research, the reliability of a binding measurement is typically assessed in a test-retest design, where PET-experiments are performed twice in a group of individuals, and the between- and within-individual variability of the measurements are evaluated^19^. Validity is the degree to which a measurement corresponds to what it is supposed to measure. A common approach to assess validity, i.e., determine whether, and the extent by which, the radioligand binds to the target of interest, is to perform a pharmacological challenge where PET measurements are conducted before and after administration of a competitor from a different chemical class.

Several such studies have been published for [^11^C]raclopride and striatum^20–22^. Extrastriatally, however, the data is sparse. Using haloperidol as a competitor Mawlawi (2001) showed that while achieving an occupancy of ~90% in striatum, only half of the purported specific binding in thalamus was displaced^21^. To our knowledge no competition experiments assessing [^11^C]raclopride binding in cortex have been published.

The aim of the present study was to explore both the validity and reliability of [^11^C]raclopride binding in extrastriatal regions. We performed a competition study in healthy controls attempting to replicate the results from Mawlawi (2001) for thalamus, but also to assess [^11^C]raclopride binding in cortex. This part of the study will from here on be referred to as COMP. In the second part, from here on referred to as TRT, we evaluated the reliability of [^11^C]raclopride binding in extrastriatal regions using a test-retest design in a separate sample of healthy controls.

## Material and Methods

### Study design

Two independent datasets were used for the competition and the test-retest design. In COMP eleven healthy male subjects (21 - 29 (25±2.5) years) participated in a previously published occupancy study of quetiapine^23^, clinical trial registration number: NCT00832221 (http://www.clinicaltrials.gov/). Quetiapine is a multimodal drug with D_2/3_R antagonist properties (K_i_ = 245 nM)^24^. Extended release (XR) or immediate release (IR) quetiapine was given once-daily during 12 days. After 4 days of dose titration of quetiapine XR from 50 mg to 300 mg, each subject received 300 mg quetiapine XR for 4 days. Treatment was then directly switched to 300 mg quetiapine IR for 4 days. The subjects participated in five PET measurements with [^11^C]raclopride: at baseline and at time for expected peak (T_max_) and trough (T_min_) plasma concentration for both drug formulations. The PET-experiments at T_max_ were performed on the fourth day of administration of XR and IR respectively and the T_min_ examination the morning after the last dose of each formulation. See the original publication for details^23^.

TRT consist of data from nine (six females) healthy subjects (37 - 71 (53±12) years) not previously published. The subjects participated in two PET measurements with [^11^C]raclopride. Time between measurements was 14 to 27 days (20±5, mean±SD). All subjects in both studies were healthy according to a clinical interview of medical history; physical examination; psychiatric interview; blood and urine chemistry; and magnetic resonance imaging (MRI) of the brain. The procedures in both studies were approved by the Research Ethics Committee in Stockholm, Sweden, and the Radiation Safety Committee at Karolinska University Hospital, Stockholm, and were performed in accordance with the 2004 revision of the Declaration of Helsinki. All subjects gave their written informed consent before participation.

### MRI

T1-weighted MRI images were acquired using a 1.5 T (COMP) or a 3 T (TRT) GE Signa system (GE Medical Systems, USA).

### Regions of interest

FreeSurfer (version 6.0, http://surfer.nmr.mgh.harvard.edu/)^25^ was used to define ten regions of interest (ROIs) on the T1-weighted MRIs of all subjects (Figure 1). ROIs were chosen based on their relevance for both neurological and psychiatric disorders, as well as for comparison with previous test-retest studies on extrastriatal [^11^C]raclopride binding^11,12^.

**FIGURE 1.**
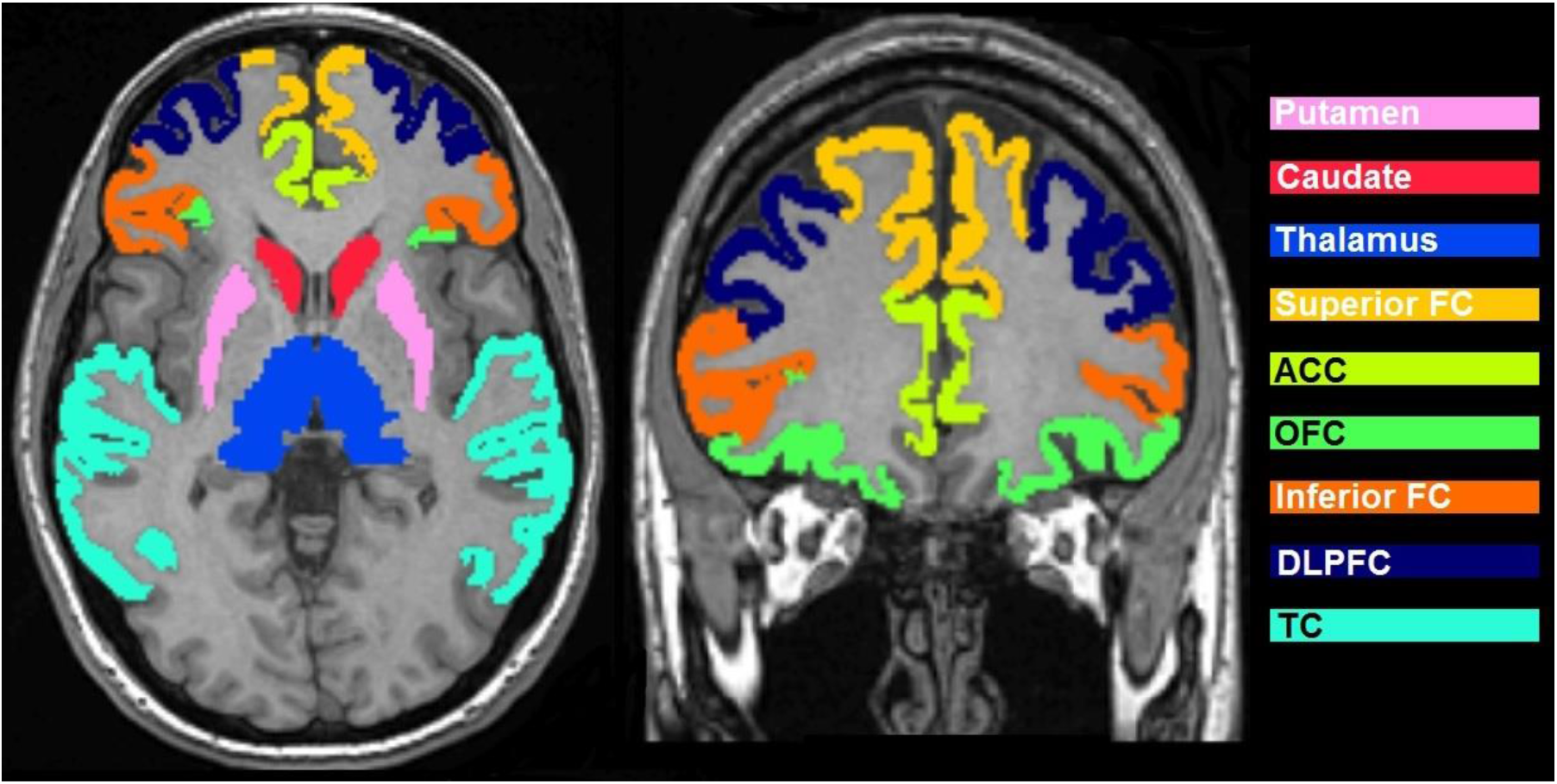
MRI for one subject from the COMP data with regions of interest overlaid. Nucleus accumbens not visible. ACC, Anterior cingulate cortex; DLPFC, dorsolateral prefrontal cortex; FC, frontal cortex; OFC, orbitofrontal cortex; TC, temporal cortex.

### Radiochemistry

[^11^C]raclopride was prepared as described previously^26^. The injected radioactivity in COMP ranged between 227-235 MBq (232±2) for the baseline examination; 207-236 (225±10) for Tmax XR (p=0.11); and 223-236 (231±5) for Tmax IR (p=0.67). The specific radioactivity was 336±264 GBq/μmol for the baseline examination; 342±280 GBq/μmol for Tmax XR (p=0.96); and 198±89 for Tmax IR (p=0.17). Injected mass was 0.32±0.17 μg for the baseline examination; 0.36±0.29 μg for Tmax XR (p=0.67); and 0.49±0.26 for Tmax IR (p=0.12). In TRT the injected radioactivity ranged between 296-524 MBq (397±98) for PET1 and 156-561 (411±135) for PET2 (p=0.80). The specific radioactivity was 148±49 GBq/μmol for PET1 and 206±75 GBq/μmol for PET2 (p=0.07) corresponding to an injected mass of 1.08±0.64 μg for PET1 and 0.82±0.52 μg for PET2 (p=0.37).

### PET experimental procedure

In each PET-experiment a saline solution containing [^11^C]raclopride was injected into a antecubital vein as a bolus (<10s). The cannula was then immediately flushed with 10 mL saline.

All subjects were examined using a high-resolution research tomograph (HRRT; Siemens Molecular Imaging, USA) with a maximum spatial resolution of ~2mm full-width-half-maximum^27^. Transmission scans were performed prior to each PET measurement in order to correct for signal attenuation.

Brain radioactivity was measured continuously, in COMP for 63 minutes and in TRT for 51 minutes. The radioactivity was reconstructed in consecutive time frames, in COMP, four 15 s, four 30 s, six 1 min, six 3 min and six 6 min frames. In TRT the initial frame sequence was identical to COMP whereas the number of 6 min frames at end of data acquisition was reduced to four.

### Quantitative analysis

PET images were corrected for head motion using a frame-to-first-minute realignment procedure^28^. Using SPM5 (Wellcome Department of Cognitive Neurology, University College, London, UK), the T1-weighted MR-images were co-registered to a summed PET-image. To obtain regional time-activity curves, the ROIs were projected onto the realigned dynamic PET-image.

From the time-activity curves, *BP_ND_* was estimated using the simplified reference tissue model (SRTM)^29^. Cerebellum, a region where specific binding has been considered negligible^8^, was used as reference. The cerebellar cortex volume was first defined using FreeSurfer, then trimmed in an automated process to include only voxels above lowest plane of pons; behind and below the posterior tip of the 4th ventricle. Only voxels located laterally of the left- and rightmost point of the 4^th^ ventricle was included. The outer layer of the resulting mask was then eroded by one voxel (Suplementary Figure S1).

### Calculations and statistics

Statistical analyses and data visualization were performed using R (version 3.3.3). Occupancy (%) of quetiapine was calculated according to the equation:

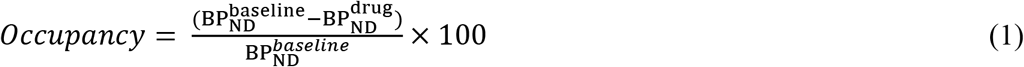

The validity of extrastriatal [^11^C]raclopride *BP_ND_* was tested comparing the baseline examination with examinations after pretreatment with quetiapine XR and IR respectively. Specific binding was defined as present when a significant (p < 0.05) decrease was showed using paired one sided t test.

Test-retest reproducibility for the TRT data was assessed using the following metrics:

*Absolute variability (VAR):*

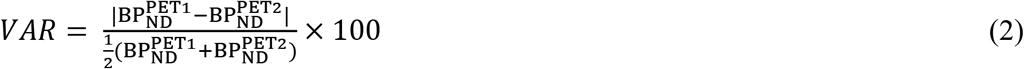

VAR is a measure of the absolute reliability of a measurement expressed as a percentage of the average *BP_ND_* value. PET1 refers to the first PET measurement, and PET2 refers to the second PET measurement. The reported value is the average VAR for all subjects.

*Intraclass correlation coefficient (ICC):*

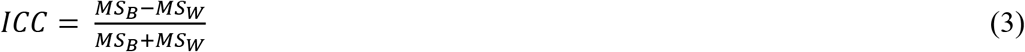

where MS_B_ denote the between subjects mean sum of squared variance and MS_W_ the within subject mean sum of squared variance. ICC normalizes the measurement error to the between-subject variance and will give information on how well a test can distinguish between individuals. The score can vary between −1 and 1, values closer to 1 indicate that most of the variance is due to between-subject rather than within-subject variation^30^.

*Standard error of measurement (SEM):*

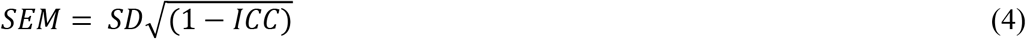

SEM is expressed in the same unit as the outcome (in this study *BP_ND_*). It is an estimate of the standard deviation of the measurement error and can be viewed as the uncertainty surrounding the outcome in a single examination^30^. Notably, though similarly named, the standard error of the *mean* and standard error of *measurement* are diverse statistical concepts.

## Results

In the COMP dataset, two subjects were excluded before image analysis due to excessive head movement during the baseline measurement. Excessive head movement was defined as more than 3 mm displacement from the reference position in more than 10% of the frames as seen in the realignment plot. In addition, PET acquisition data from the quetiapine IR measurement for one subject was excluded due to a delay of the examination of two hours beyond expected T_max_ for the plasma concentration of the drug. Nine subjects were included in the final analysis of baseline and quetiapine XR data. For the quetiapine IR data eight subjects were analyzed. In the TRT dataset SRTM failed in the anterior cingulate cortex in one individual producing a negative *BP_ND_* value. This value was excluded from further analysis.

Results from COMP are shown in Figure 2. In extrastriatal regions a significant decrease of *BP_ND_* was seen only in thalamus and temporal cortex (TC) after treatment with XR as well as IR formulations of quetiapine (Table 1). In putamen the occupancy was 33±11% and 51±4% (mean±SD) in the quetiapine XR and IR measurements respectively. Occupancy was lower in extrastriatal regions: 10±14% and 20±17% in thalamus and 12±11% and 18±17% in TC (Table 1).

**FIGURE 2.**
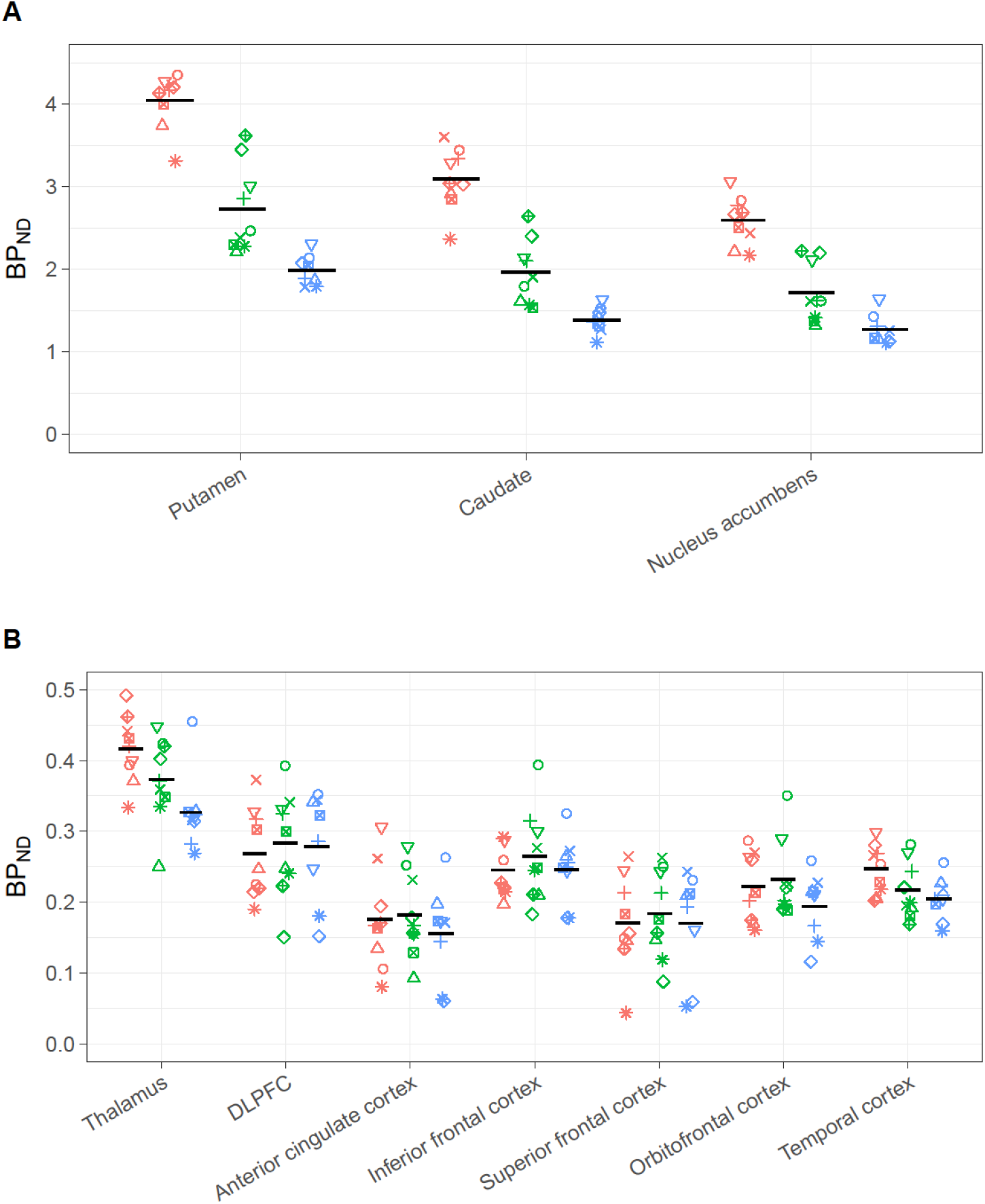
COMP [^11^C]raclopride binding data. A) Striatal ROIs (for reference). Each ROI represents three PET examinations, from left to right: Baseline (red); at T_max_ post quetiapine XR (green); at T_max_ post quetiapine IR (blue). Horizontal bars represent mean *BP_ND_*. B) Extrastriatal ROIs, same order of PET examinations as in A. DLPFC, dorsolateral prefrontal cortex.

**TABLE 1.** Quetiapine occupancy data

| Region | Baseline (n = 9) | Quetiapine XR (n = 9) |  |  | Quetiapine IR (n = 8) |  |  |
| --- | --- | --- | --- | --- | --- | --- | --- |
| | Mean ± SD<br>(BP<sub>ND</sub>) | Mean ± SD<br>(BP<sub>ND</sub>) | test vs<br>baseline<br>(p) | occ<br>(%) | Mean ± SD<br>(BP<sub>ND</sub>) | test vs<br>baseline<br>(p) | occ (%) |
| Putamen | 4.05±0.33 | 2.73±0.53 | <0.001 | 32.58 | 1.99±0.19 | <0.001 | 50.62 |
| Caudate | 3.1±0.37 | 1.97±0.39 | <0.001 | 36.14 | 1.39±0.16 | <0.001 | 54.96 |
| Nucleus Accumbens | 2.6±0.29 | 1.72±0.36 | <0.001 | 33.86 | 1.27±0.18 | <0.001 | 50.75 |
| Thalamus | 0.42±0.05 | 0.37±0.06 | 0.029 | 9.93 | 0.33±0.06 | 0.007 | 19.53 |
| DLPFC | 0.27±0.06 | 0.28±0.07 | 0.747 | -7.49 | 0.28±0.08 | 0.556 | -3.19 |
| Anterior cingulate | 0.18±0.07 | 0.18±0.06 | 0.616 | -15.82 | 0.16±0.07 | 0.287 | -2.48 |
| Inferior frontal cortex | 0.25±0.04 | 0.26±0.07 | 0.865 | -7.7 | 0.25±0.05 | 0.465 | -0.44 |
| Superior frontal cortex | 0.17±0.07 | 0.18±0.06 | 0.782 | -23.26 | 0.17±0.07 | 0.418 | -2.67 |
| OFC | 0.22±0.05 | 0.23±0.06 | 0.602 | -2.37 | 0.19±0.05 | 0.059 | 12.63 |
| Temporal cortex | 0.25±0.03 | 0.22±0.04 | 0.007 | 12.07 | 0.2±0.03 | 0.01 | 17.75 |

P-values calculated using one sided paired t tests. BP<sub>ND</sub>, binding potential using simplified reference tissue model; DLPFC, dorsolateral prefrontal cortex; occ, occupancy; OFC, orbitofrontal cortex; SD, standard deviation

TRT was completed in nine control subjects. ICC values were higher and VAR values were lower in striatal ROIs, compared to extrastriatal regions (Table 2).

**TABLE 2.**
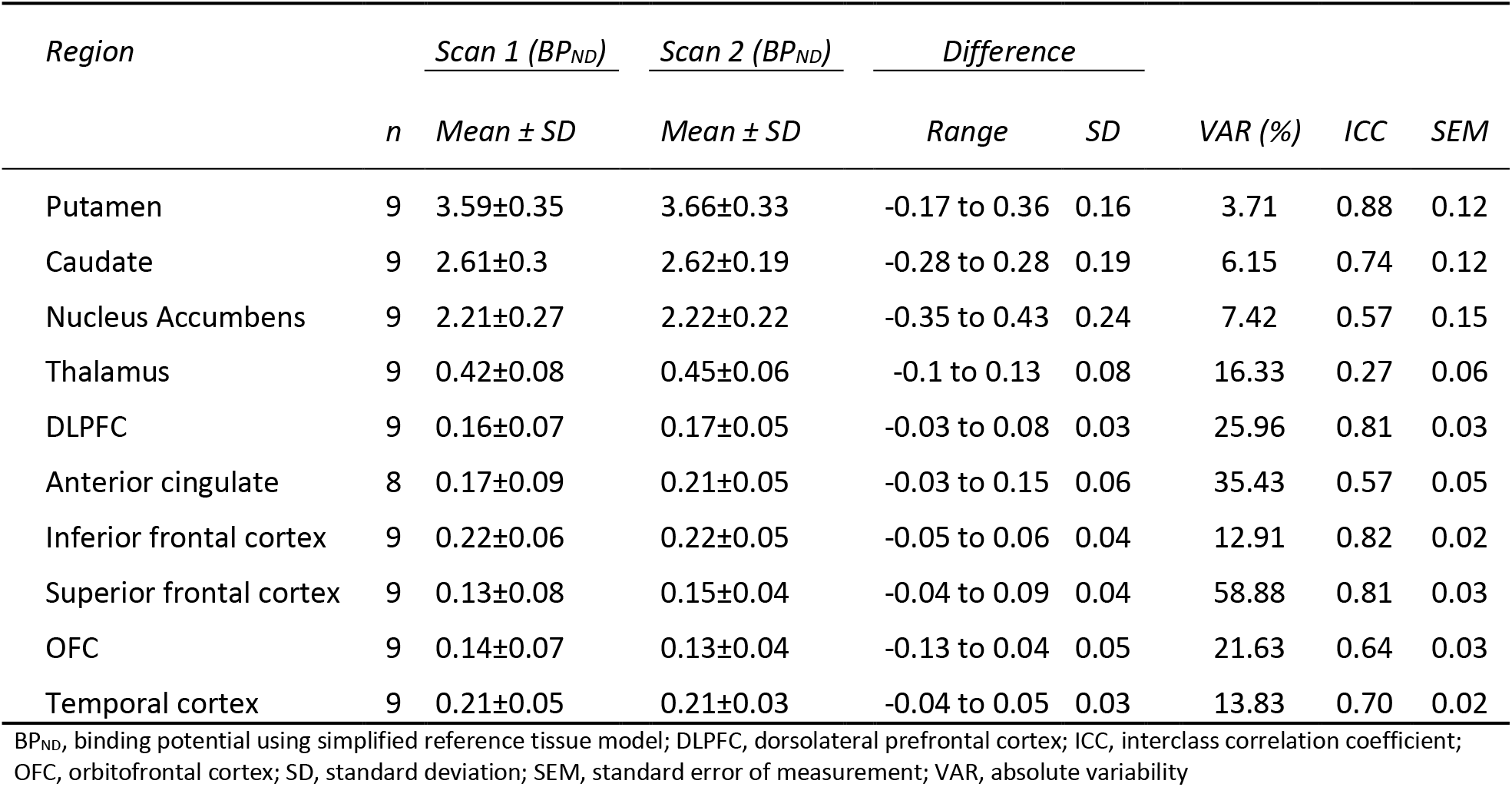
[^11^C]raclopride test-retest data; BP_ND_ values and statistics

## Discussion

We have examined the validity and reliability of the extrastriatal binding characteristics of [^11^C]raclopride. No specific raclopride binding could be detected in most examined cortical areas, as determined using a pharmacological competition analysis. In the thalamus and TC ROIs we observed some indication of specific binding, although occupancy was lower than in striatum. Further, the test-retest repeatability of extrastriatal *BP_ND_* was low in our data.

Our results have implications when interpreting and planning clinical studies. Similar to Mawlawi (2001) the COMP data indicate that only half of the calculated *BP_ND_* in thalamus reflect specific binding. Estimations of effect sizes need to be adjusted accordingly. Assuming that 10% difference in the density of D_2/3_R in thalamus is considered a relevant clinical finding in a cross-sectional study, the corresponding apparent effect size would be ~0.8, requiring 25 subjects per group for 80% power. Our results however indicate that the actual effect size would be ~0.3, translating into 175 subjects per group. Importantly, a non-significant finding in an extrastriatal region in a [^11^C]raclopride study powered for striatal regions will give very little information on whether an effect is present or not.

We investigated the test-retest repeatability of [^11^C]raclopride binding in a sample with a clinically relevant age- and gender diversity. We observed a VAR of 3.7-7.4% and ICC between 0.57-0.88 in striatal regions. The results are similar to a previous test-retest study of [^11^C]raclopride in a bolus-constant infusion protocol and a high resolution PET system^11^, and numerically superior to previously published lower resolution PET data^31,32^. The reliability of our data was lower in extrastriatal regions compared to striatum (Table 2). We were not able to replicate the VAR values of 3.7-13.1% or ICC of 0.64-0.92 in the extrastriatal regions reported by Alakurtti et al (2015). Further, it should be noted that before validity is proven it is difficult to interpret ICC and VAR, or rather: poor values still indicate a problem even if the validity is good, but before accepting a high ICC or low VAR as indicative of reliable *specific* binding, validity need to be established.

Our data indicate that the greater part of [^11^C]raclopride *BP_ND_* measured in neocortex does not reflect specific binding. However, since we consistently measure higher [^11^C]raclopride signal in, e.g., frontal cortex, compared to cerebellum the question arises to what this difference should be ascribed if not to specific binding? The explanation suggested by Mawlawi (2001) is a systematically lower non-displaceable compartment (*V_ND_*) in cerebellum compared to cerebral target regions^21^, a *V_ND_*-bias. This interpretation is in line with our observations of lower occupancy in regions with lower densities of D_2/3_R (see Figure S2 for an explanation on how *V_ND_*-bias propagates to occupancy values). The presence of a discrete difference in *V_ND_* between target and reference will not matter much in receptor rich regions (i.e. striatum) but will become a serious validity issue in low-binding regions. If, for example, *V_ND_* is 10% lower in the reference region then the “true” *BP_ND_* in the target region will be falsely increased with 0.1 and 10%^33^. In, e.g., frontal cortex where we might have a “true” [^11^C]raclopride *BP_ND_* of 0.05 or less, even a small *V_ND_*-bias would thus be highly problematic. However, since the protocol did not include arterial blood sampling, a more detailed analysis of *V_ND_* in different ROIs was not possible.

There are other possible explanations for the observed differences in quetiapine occupancy between high and low density D_2/3_R regions: (i) quetiapine could have different occupancy in different brain regions. In the time span between baseline- and post drug examinations quetiapine could (ii) cause the extrastriatal expression of D_2/3_R to increase, or (iii) cause the concentration of endogenous dopamine to decrease. However, several previous occupancy studies of quetiapine at steady-state using high affinity radioligands have shown similar or higher occupancy of D_2/3_R in cortex compared to striatum^34,35^ and no study has, to our knowledge, shown lower occupancy. This makes i-iii unlikely explanations to our findings.

There are some limitations to this study. The standardized uptake value (SUV) in cerebellum was lower in the examinations performed after pretreatment with quetiapine compared to baseline (supplement, Figure S3 and Table S1). This may be explained by (i) presence of specific [^11^C]raclopride binding to D_2/3_R in cerebellum; (ii) that quetiapine displaces non-specific binding of raclopride, or (iii) that quetiapine decreases [^11^C]raclopride brain uptake. (i) will result in an underestimation of occupancy equally in low- and high binding regions and would thus not alter the conclusions of our results^33,36^. The same is true for (ii) given that the displacement of non-specific binding is equal in all regions. Additionally, we observed that centrum semiovale (Figure S1), a region containing only white matter, showed similar decrease of SUV (Figure S3) which lends support to explanation (ii) and (iii). The explanation we find most probable, (iii), would also likely not affect our results since the decrease of measured radioactivity would be proportional in target and reference regions.

Regarding the test-retest dataset, a caveat that should be highlighted is the fact that time between examinations was 20±5 days. Most commonly, PET test-retest examinations are performed within 1-2 days. This protocol was chosen to mimic that of typical clinical studies where patients are examined repeatedly under an extended period of time, an established test-retest design for evaluation of clinical applicability^32,37^.

## Conclusions

In most brain regions outside striatum, we could not find proof of valid [^11^C]raclopride binding quantification, as little or no decrease in *BP_ND_* was seen after administration of a competitor. Further, we found extrastriatal test-retest repeatability to be poor. While confirming the validity and reliability of [^11^C]raclopride binding quantification in striatum, our findings indicate that [^11^C]raclopride PET not is a suitable tool for D_2/3_R binding quantification in extrastriatal regions. Before validity is proven strong caution is warranted when interpreting studies applying [^11^C]raclopride for measuring of D_2/3_R availability in extrastriatal regions.

