## Supplement for "Validity and reliability of extrastriatal [^11^C]raclopride binding quantification in the living human brain"

**Supplementary material**

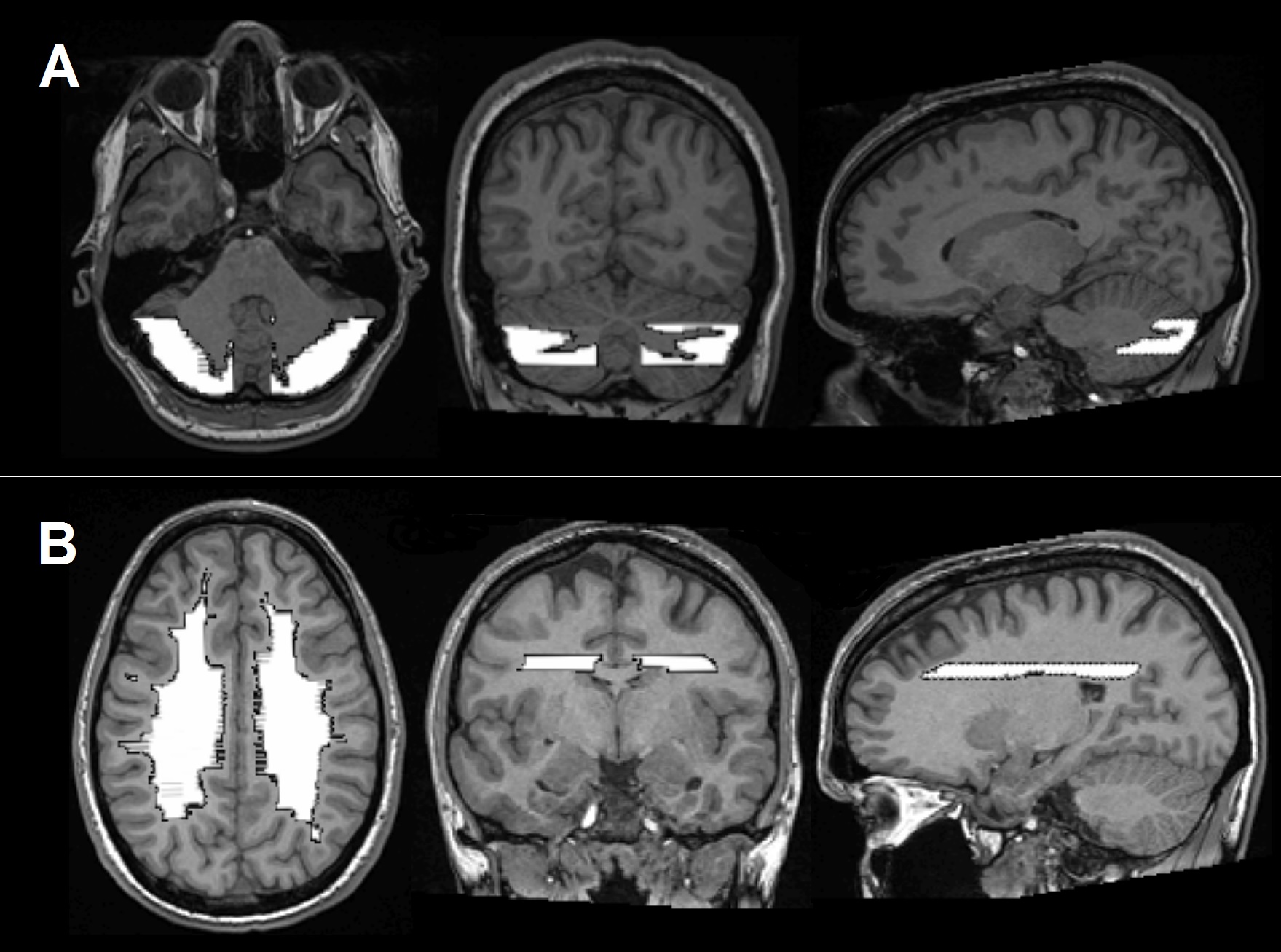

**Figure S1** A) MRI from one subject in the COMP dataset showing the delineation of the reference region used both in COMP and TRT B) Centrum semiovale in the same subject.

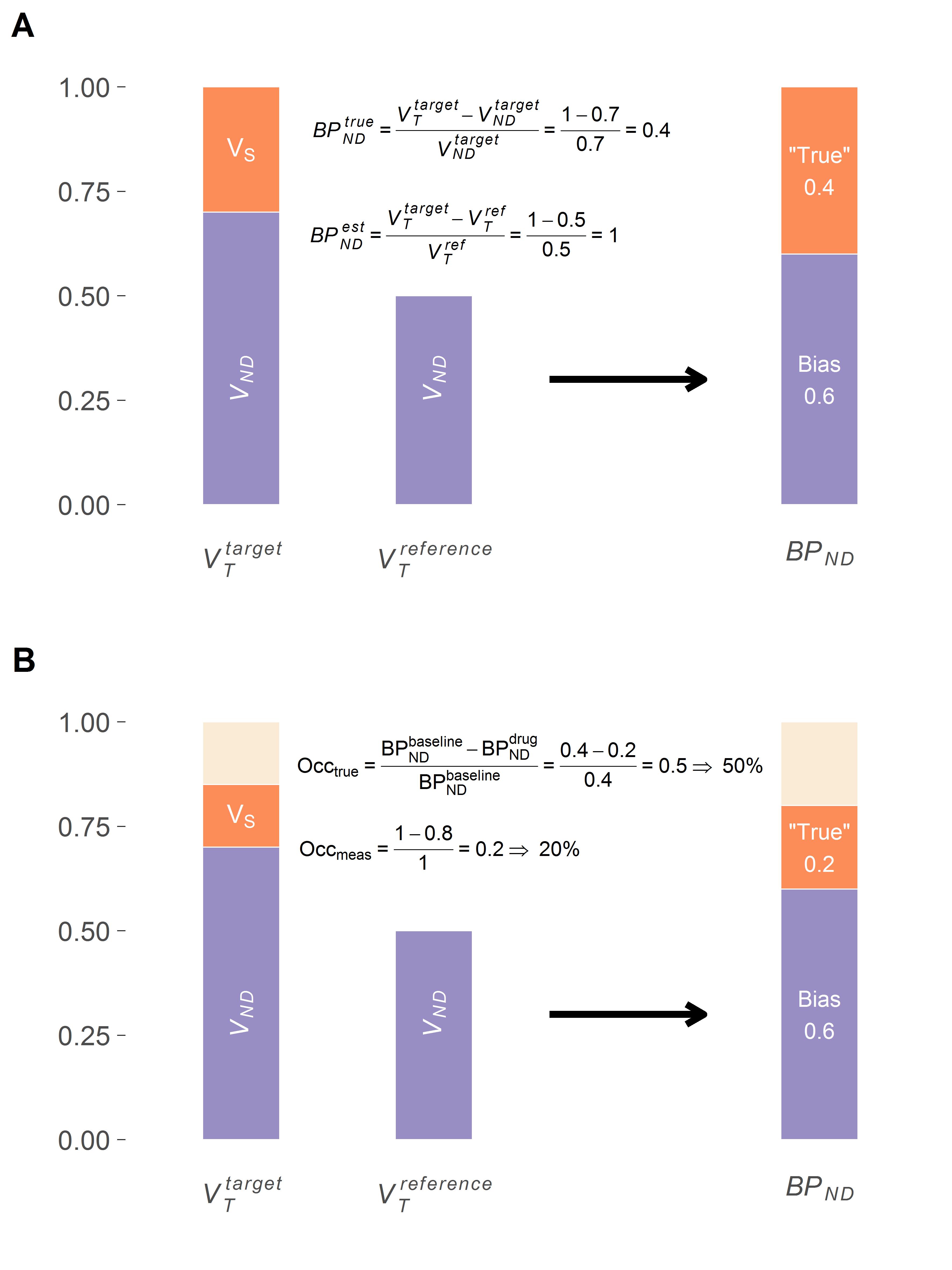

**Figure** **S2** A) In a target region with higher non-displaceable signal than in the reference region *BP_ND_* is over-estimated and occupancy under-estimated. In this example 40% higher *V_ND_* in the target region (i.e. 0.5 compared to 0.7) causes an inflated *BP_ND_* estimate in reference tissue models. B) illustrates how the same discrepancy propagates into an underestimation of occupancy (20% rather than the true 50%). Bars to left of the arrow represents distribution volumes (*V_T_*) separated in specific binding (*V_S_*) and non-displaceable signal (*V_ND_*). Leftmost bar is the target region and next to it is the reference region. (A) Baseline examination; (B) the same region after 50% of *V_S_* is blocked. *BP_ND_*, non-displaceable binding potential; Occ, occupancy

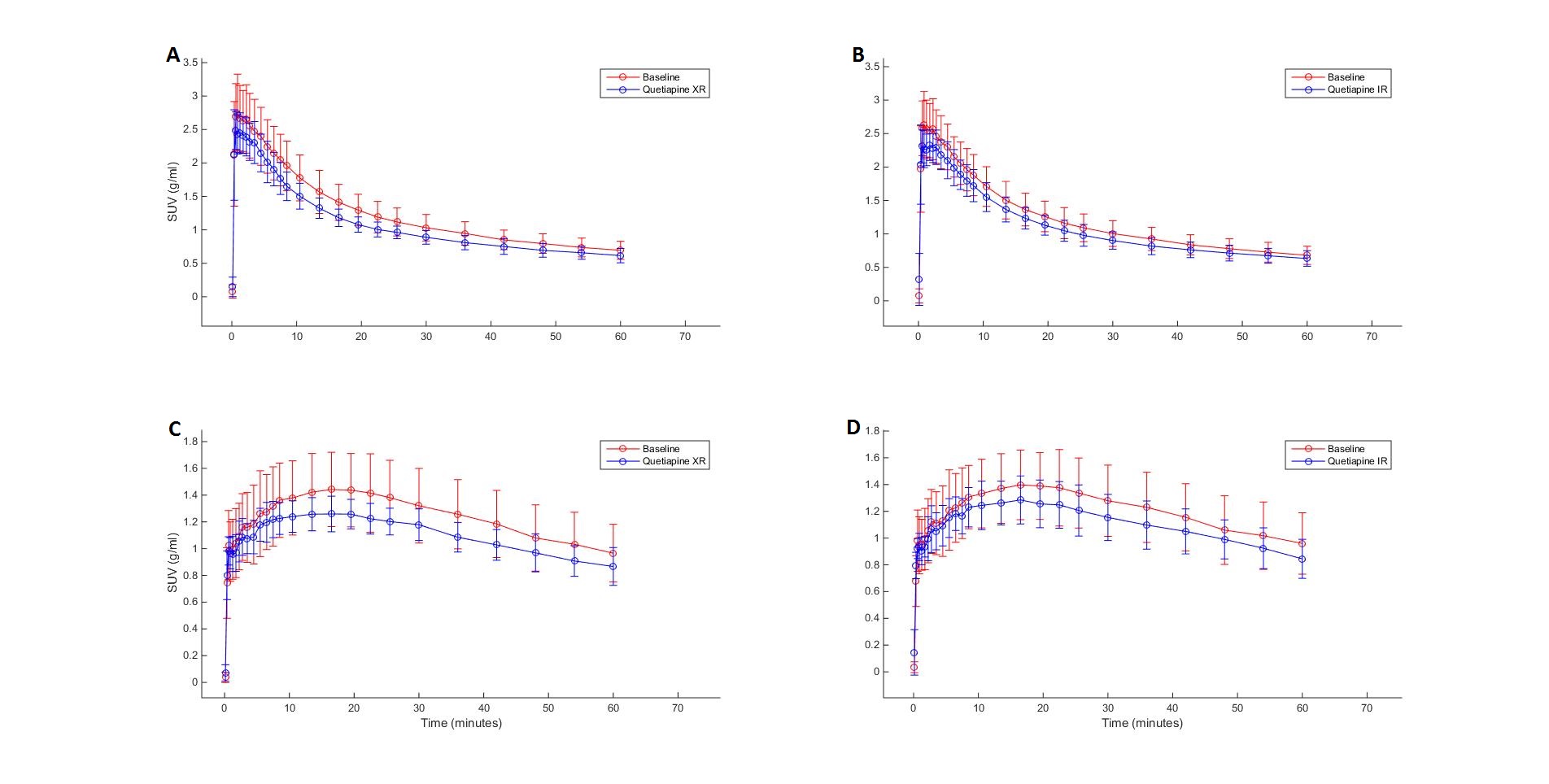

**Figure S3.** Average time activity curves (TACs) standardized to injected radioactivity and weight. **A:** TAC in gray matter cerebellum at baseline (red) and after pretreatment with quetiapine XR (blue).**B**: TAC in the same region at baseline (red) and after pretreatment with quetiapine IR (blue). **C:** TAC in centrum semiovale baseline (red) and after pretreatment with quetiapine XR (blue). **D:** TAC in the same region after pretreatment with quetiapine IR. Error bars show standard deviation. N = 9 for quetiapine XR; N = 8 for quetiapine IR.

| **Table S1.** |  |  | |  |  |  |  |  |
| --- | --- | --- | --- | --- | --- | --- | --- | --- |
| *Region* | *Quetiapine XR Tmax (n=9)* | | | |  | *Quetiapine IR Tmax (n=8)* | | |
|  | *SUV baseline* | *SUV drug* | *Test (p)* | |  | *SUV baseline* | *SUV drug* | *Test (p)* |
| Cerebellar gray | 37.2±6.9 | 32.2±4.1 | <0.01 | |  | 36.4±6.9 | 32.9±5 | 0.02 |
| Centrum semiovale | 48.7±10.2 | 42.8±4.4 | 0.04 | |  | 47.6±10.3 | 43.1±6.6 | 0.12 |
| Standardized uptake values (SUV) calculated as area under the curve from frame 18 to the end (18-63 min)  for time activity curves standardized to injected radioactivity and weight. Baseline tested against drug  using paired t-test. | | | | | | | | |
